# Evolution of sex chromosomes is prior to speciation in the dioecious *Phoenix* species: Sex chromosome evolution in *Phoenix*

**DOI:** 10.1101/033365

**Authors:** Emira Cherif, Salwa Zehdi-Azouzi, Amandine Crabos, Karina Castillo, Nathalie Chabrillange, Jean-Christophe Pintaud, Amel Salhi-Hannachi, Sylvain Glémin, Frédérique Aberlenc-Bertossi

**Affiliations:** IRD/CIRAD F2F-palmiers Group, UMR DIADE, Centre IRD, 911 avenue Agropolis, 34394, Montpellier, France; Laboratoire de Génétique Moléculaire, Immunologie et Biotechnologie, Faculté des sciences de Tunis, Université Tunis El Manar, Campus Universitaire, 2092, El Manar, Tunisia; Dynadiv, UMR DIADE, Centre IRD, 911 avenue Agropolis, 34394, Montpellier, France; Institut des Sciences de l’Evolution de Montpellier, Unité Mixte de Recherche 5554 (Université de Montpellier-CNRS-IRD-EPHE), 34095, Montpellier, France; Department of Ecology and Genetics, Evolutionary Biology Centre, Uppsala University, SE-752 36 Uppsala, Sweden.

**Keywords:** sex-linked gene, sex chromosomes, recombination arrest, dioecy, speciation

## Abstract

Understanding the driving forces and molecular processes underlying dioecy and sex chromosome evolution, leading from hermaphroditism to the occurrence of male and female individuals, is of considerable interest in fundamental and applied research. The genus *Phoenix*, belonging to the family Arecaceae, consists of only dioecious species. Phylogenetic data suggests that the genus *Phoenix* diverged from a hermaphroditic ancestor shared with its closest relatives. Here we investigated the evolution of suppressed recombination within the genus *Phoenix* as a whole by extending the analysis of *P*. *dactylifera* sex-related loci to eight other species within the genus. We also performed a phylogenetic analysis of a date palm sex-linked *PdMYB1* gene in these species. We found that X and Y sex-linked alleles clustered in a species-independent fashion. Our data show that sex chromosomes evolved before the diversification of the extant dioecious species. Furthermore, the distribution of Y haplotypes revealed two male ancestral paternal lineages which may have emerged prior to speciation.

## 1. Introduction

Dioecy, in angiosperms, may result from two sex-determining mutations, a recessive male-sterility mutation and a mutation at a linked locus causing the loss of female functions [1]. If both these partially linked mutations establish polymorphisms [1, 2], closer linkage between the two sex-determination loci is favored by selection, to maintain the correct combination of mutations, and avoid sterile recombinants. This may explain the suppressed recombination that characterizes sex chromosomes. The time since recombination stopped in the sex-determining region defines the age of the sex chromosome system, and it is also of interest to know whether a single recombination suppression occurred, or multiple events, resulting in several evolutionary strata having evolved, as has occurred in mammals, *Silene latifolia*, and *Papaya carica* [3, 4]. Dioecy and sex chromosomes have evolved repeatedly and independently in different plant taxa [5, 6, 7]. However, only a few systems have been described in detail. Understanding sex chromosome emergence during the evolution of dioecy, leading from hermaphroditism to the occurrence of male and female sterile individuals, is therefore of major fundamental interest, with many potential agronomic applications.

The genus *Phoenix* (Arecaceae, Coryphoideae, Phoeniceae) includes fourteen dioecious species, distributed from the Atlantic islands throughout the Mediterranean region, Africa, Middle East, and as far as southern Asia to the northwestern Pacific [8,9,10].

DNA sequence divergence from other palm genera is high [11, 12], and it has been suggested that the genus *Phoenix* might possess an ancient sex chromosome system [20]. The tribe Phoeniceae is sister to the predominantly hermaphroditic tribe Trachycarpeae [13], but is distinguished by several morphological differences [10], and the divergence time is estimated to be around 49 ± 16 mya [14]. Assuming a single origin of dioecy in *Phoenix*, this date gives an upper bound to the age of the sex-linked non-recombining region. Interspecific relationships within *Phoenix* were studied by Pintaud *et al*. [15] based on two chloroplast loci (psbZ-trnfM and rpl16-rps3), recovering five phylogenetic lineages, namely *P. loureiroi-acaulis-pusilla*, *P. roebelenii-paludosa*, *P. caespitosa*, *P. reclinata* and a larger lineage consisting of *P. dactylifera, P. atlantica, P. theophrasti, P. sylvestris* and *P. rupicola*. Sexual dimorphism in the genus *Phoenix* has been dated back to the Eocene period (between 33.9 and 55.8 million years ago) [10, 16] on the basis of fossil records of *Phoenix* male flowers. Dioecy could thus be very ancient within the genus. In *P*. *dactylifera*, sex differentiation results from the arrest of male or female organ development in the initial bisexual flower buds [17] and the species has an XY sex chromosome system [18,19,20]. A non-recombining XY-like region was inferred in the date palm genome, based on 3 microsatellite loci showing alleles confined to males, and two different Y haplogroups were found [20]. Recently, Mathew *et al*. [21] constructed a genetic map of date palm and localized the sex segregating region to LG12. The physical length of this region is estimated to be 13Mb, about 2% of the genome [21].

All known species in the genus *Phoenix* are dioecious. Dioecy is probably an ancestral character in the genus. However, it is important to test explicitly whether sex chromosomes evolved before speciation within the genus, to exclude the possibility that dioecy evolved in more than one lineage, and to test whether suppressed recombination might have evolved in only certain lineages. If X and Y alleles of different species cluster together, rather than by clustering within their respective species, sex linkage must have evolved before speciation. Conversely, if these alleles cluster according to species, then sex linkage must have evolved after speciation [22].

We used sex-linked markers identified in *P. dactylifera* [20] to study eight other species, and found that one sex-linked MYB gene, PdMYB1, was present in seven of the studied species. Our results provide strong evidence that sex evolved before the appearance of the extant species of the genus *Phoenix*.

## 2. Material and Methods

### (a) Plant material

Nine of the 14 *Phoenix* species were studied, mainly from natural populations (including 64 males and 70 females). Three species with large samples in our study are widespread: *Phoenix dactylifera* (34), *P. reclinata* (10), *P. sylvestris* (18), while three species have restricted distribution: *P. atlantica* (17), *P. canariensis* (21) and *P. roebelenii* (24) were included (Figure 1), and three other species only small samples, *P. acaulis*, *P. rupicola* (one male and one female) and *P. loureroi* (three males and three females) (Figure 1); these three species were excluded from the statistical analyses (electronic supplementary material, Table 1).

**Figure 1.**
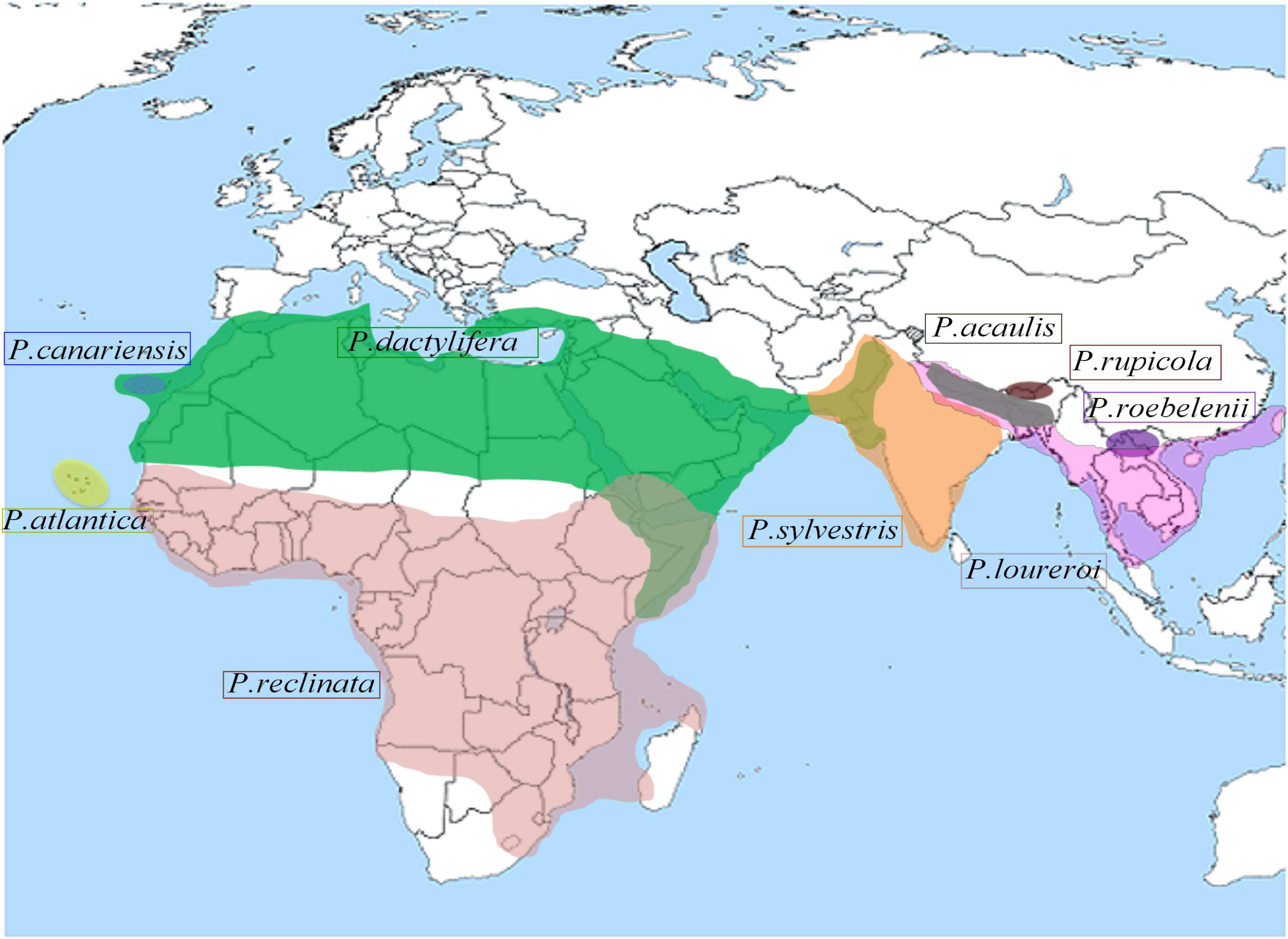
Geographical distribution of the studied *Phoenix* species.

### (b) DNA extraction

Leaf samples were freeze-dried for 72 h with an Alpha1–4LD Plus lyophilizer (Fisher Scientific, Illkirch, France) and ground with a Tissue Lyser System (Qiagen). DNA extraction was carried out using the Dneasy plant mini kit (Qiagen) according to the manufacturer’s instructions. All samples were adjusted to 10 ng.µl^−1^ concentration for subsequent analyses.

### (c) PCR assays

To test the transferability of date palm sex-linked markers to the other *Phoenix* species, the three SSR loci (mPdIRD50, mPdIRD52 and mPdIRD80) designed by Cherif *et al*. [20] were tested across eight newly studied species and new genotypes of *P.dactylifera*.

PCR reactions were performed in an Eppendorf thermocycler (AG, Hamburg, Germany). The reaction volume was 20 µl and contained 10 ng of genomic DNA, 2 µl of 10X reaction buffer with 2 mM MgCl_2_, 200 µM dNTPs, 1 U Taq polymerase, 10 pmol of the fluorochrome-marked forward primer and of the reverse primer and MilliQ water. The PCR reactions were carried out with following parameters: 5 min denaturation at 95°C, followed by 35 cycles of 95°C for 30 s, 56°C for 30 s and 72°C for 45 s, and then a final elongation step at 72°C for 10 min. PCR products were analysed using an ABI 3130XL Genetic Analyzer (Applied BioSystems, Foster City, CA, USA). Allele sizes were scored with GeneMapper software v4.0 (Applied BioSystems).

### (d) Cloning and sequencing of *PdMYB1* sequences

In order to verify that the size conservation of sex-linked SSR across *Phoenix* species is not due to size homoplasy, the mPdIRD80 locus was sequenced. In addition, potential genes around the three date palm sex-linked markers [20] were searched in the published genome assembly [19]. A *MYB* gene called *PdMYB1* was identified by BLASTn on GenBank database near the mPdIRD80 locus.

Primers PdMYB1R1 or PdMYB1R2 and PdMYB1F5 or mPdIRD80F (electronic supplementary material, Table 2) were used to amplify the target *MYB* gene from seven *Phoenix* species (*P.dactylifera, P.reclinata, P.atlantica, P.sylvestris, P.roebelenii, P.rupicola, P.canariensis*) (electronic supplementary material, Figure B). All sequences were amplified from genomic DNA, using Taq polymerase (GoTaq G2 DNA Polymerase, Promega). Reactions were performed in 20µL, with following final concentrations: 1X Green Buffer, dNTPs 0.2mM each, primers 0.5µM each, GoTaq 1U/tube and 50ng DNA/tube. The PCR conditions were as follows: one incubation at 95°C for 2 min; 30 cycles of: denaturing at 95°C for 30s, annealing at 57°C for 30s, and elongation at 72°C for 2 min; and a final extension at 72°C for 5 min. PCR products were separated on 1% agarose gel and stained with ethidium bromide. PCR products matching the target size under UV light were purified using a PCR Clean-Up Kit (Wizard SV Gel and PCR Clean-Up System, Promega) and cloned into pGEM-T Easy vector (pGEM-T Easy Vector System I, Promega) according to the manufacturer’s conditions. The ligation products were transformed into JM109-competent cells (Bacterial Strain JM109, Glycerol Stock, Promega) and positive colonies were confirmed by PCR using previous primers. Plasmids of positive colonies were isolated (Wizard Plus SV Minipreps DNA Purification System Promega) and adjusted to a concentration of 75ng/µl for sequencing using SP6 and T7 primers at Eurofins Genomics (Germany). Sequence results in fasta format were assembled and analyzed using SeqMan Pro software (Lasergene). X and Y alleles were respectively obtained from homozygous XX female plants and from XY male plants. To identify western and eastern Y alleles, western and eastern male alleles of *P. dactylifera* [20] were used as a reference.

### (e) Genetic analyses

Much of our analysis used the GenAlEx 6.5 program [23, 24], including analyses of allele frequencies and allele size distributions in male and female individuals within *Phoenix* species, estimates of genetic variability, including values of Ho, He, F*is* and R*_st_* values used to characterize subdivision between males and females (electronic supplementary material, Table 3, Table A, Table 4 and 5).

For the remainder of the study, each *Phoenix* species was considered as a population and the subsequent genetic analyses focused on male and female, treated as separate groups in tests for subdivision and differences in heterozygote frequencies. R*_st_* was estimated according to the stepwise mutation model [25] (Table 1), and the AMOVA procedure implemented in GenAlEx 6.5 [23, 24] used microsatellite distance matrix data; significance was determined by running 10 000 permutations (electronic supplementary material, Figure 1). Principal component analysis (PCA) (electronic supplementary material, Table A) was performed using the *dudi.pca* function, implemented in ade4 package for the R software v2.15.3 [26] (electronic supplementary material, Figure A).

**Table 1.**
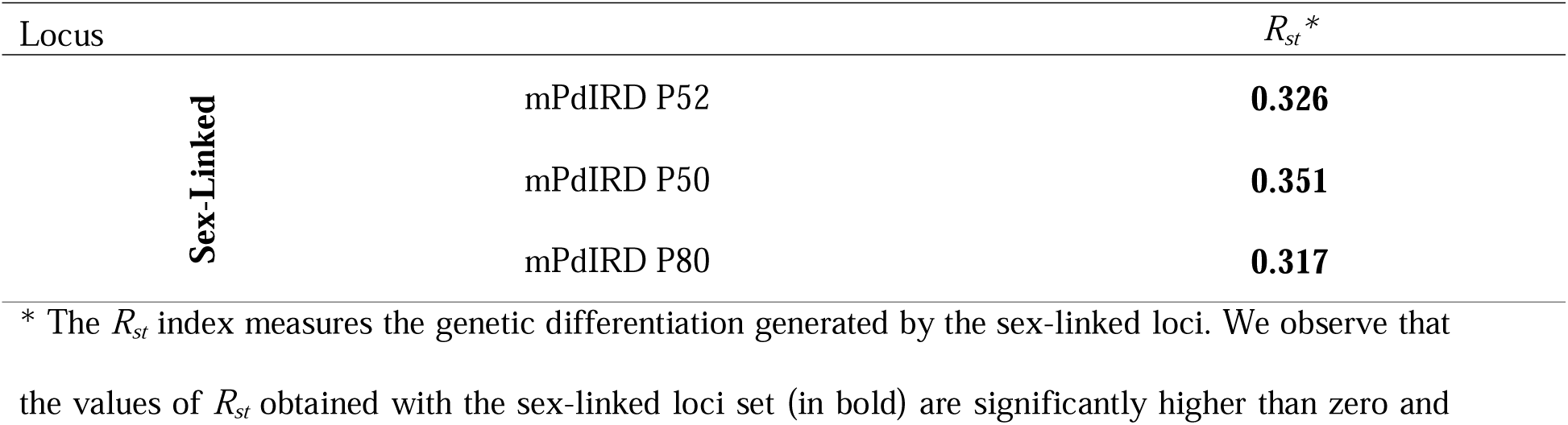
Genetic structuring within the studied *Phoenix* species generated by each sex-linked microsatellite between the two sexes.

Expected heterozygote frequencies assuming Hardy–Weinberg genotype frequencies (He) were compared with those observed (Ho) within males and females of *Phoenix* species using the GenAlEx 6.5 program [23, 24].

Two different *Fis* estimates, namely Weir and Cockerham’s estimate [27] and Robertson and Hill’s estimate [28] (electronic supplementary material, Table 5), were used to test whether there was an excess of heterozygotes in the two groups (male and female) within each *Phoenix* species. The second estimate has a lower variance under the null hypothesis [28]. To determine the significance of *Fis*, P-values were calculated by the Markov chain method [29] (electronic supplementary material, Table 6) using the Genepop 4.2 program [30, 31].

To infer X and Y haplotypes within *Phoenix* species, the alleles of the three sex-linked loci (mPdIRDP80, mPdIRDP50 and mPdIRDP52) that were most frequently associated were determined for each genotype in each species, using the EM algorithm implemented in the PowerMarker program [32]. Additionally, the EM algorithm was used to resolve the genotype of each male or female individual into the two most probable haplotypes (electronic supplementary material, Note 1).

The Y-SSR haplotypes of the genus were used to construct a median joining haplotype network [33] using the Network 4.6.1.1 program. We assumed a single-repeat mutation model, and repeat variants were given a weighting of 10. To highlight the clustering of Y and X alleles within the genus, a multiple alignment of *PdMYB1* genomic sequences from the studied *Phoenix* species was performed using CLC Sequence Viewer v7.02 software (CLC BIO) (electronic supplementary material, Figure C). The alignment was then used as an input in MEGA6.0 [34] and a tree was produced, using coding and non-coding sequence, (gaps excluded) by a Maximum Likelihood method under the Kimura 2-parameter model [35]. The latter model was chosen after comparison of the BIC scores (Bayesian Information Criterion) between different models (electronic supplementary material, Table 7 and Figure D). The same results were obtained by other Neighbor-joining and maximum parsimony. The ortholog of *PdMYB1* in the *Elaeis guineensis* genome was used as an outgroup in phylogeny reconstruction.

## 3. Results and Discussion

*Phoenix* species were chosen to represent the main evolutionary lineages on the basis of previously established morphological, anatomical and molecular phylogenies [8, 15, 36]. The geographical distribution of the nine *Phoenix* species analyzed is shown in Figure 1 [8, 9, 10]. The species studied are characterized by a wide and continuous distribution from the Canary and Cape Verde islands in the Atlantic Ocean, reaching as far as Taiwan through Africa, Madagascar, the Middle East, Pakistan and India. *P*. *canariensis* and *P. atlantica* are respectively endemic to the Canary and the Cape Verde islands. *P. roebelenii*, *P. rupicola* and *P. acaulis* have restricted ranges, respectively limited to the north of Laos and Vietnam and the north of India and Nepal. *P. dactylifera* has the largest distribution area, which overlaps that of *P. canariensis, P. reclinata* in the Horn of Africa and *P. sylvestris* on the Indian subcontinent.

### (a) Conservation of the sex-linked loci within overall the studied *Phoenix* species

Three sex-linked SSR loci, mPdIRD50, mPdIRD52 and mPdIRD80, were identified in *P*. *dactylifera* by Cherif *et al*. [20]. We mapped mPdIRD50 and mPdIRD80 on the sex linkage group (LG12) [21]. The mPdIRD52 has not mapped on the available genetic map. The three sex-linked loci were successfully amplified in all individuals of the eight additional species analysed.

#### (i) Allele frequencies

To further test whether sex linkage is conserved between *P. dactylifera* and the other *Phoenix* species, Y- and X-linked allele frequencies for each locus were investigated.

##### mPdIRDP52 locus

The mPdIRDP52 locus yielded a total of 18 alleles. Four were Y-linked (mPdIRDP52_188, mPdIRDP52_189, mPdIRDP52_191 and mPdIRDP52_193) exclusively distributed within males. *P. roebelenii* was found to have a private Y-allele (mPdIRDP52_189) in addition to the mPdIRDP52_191 allele (Figure 2). The fourteen remaining alleles were X-linked.

**Figure 2.**
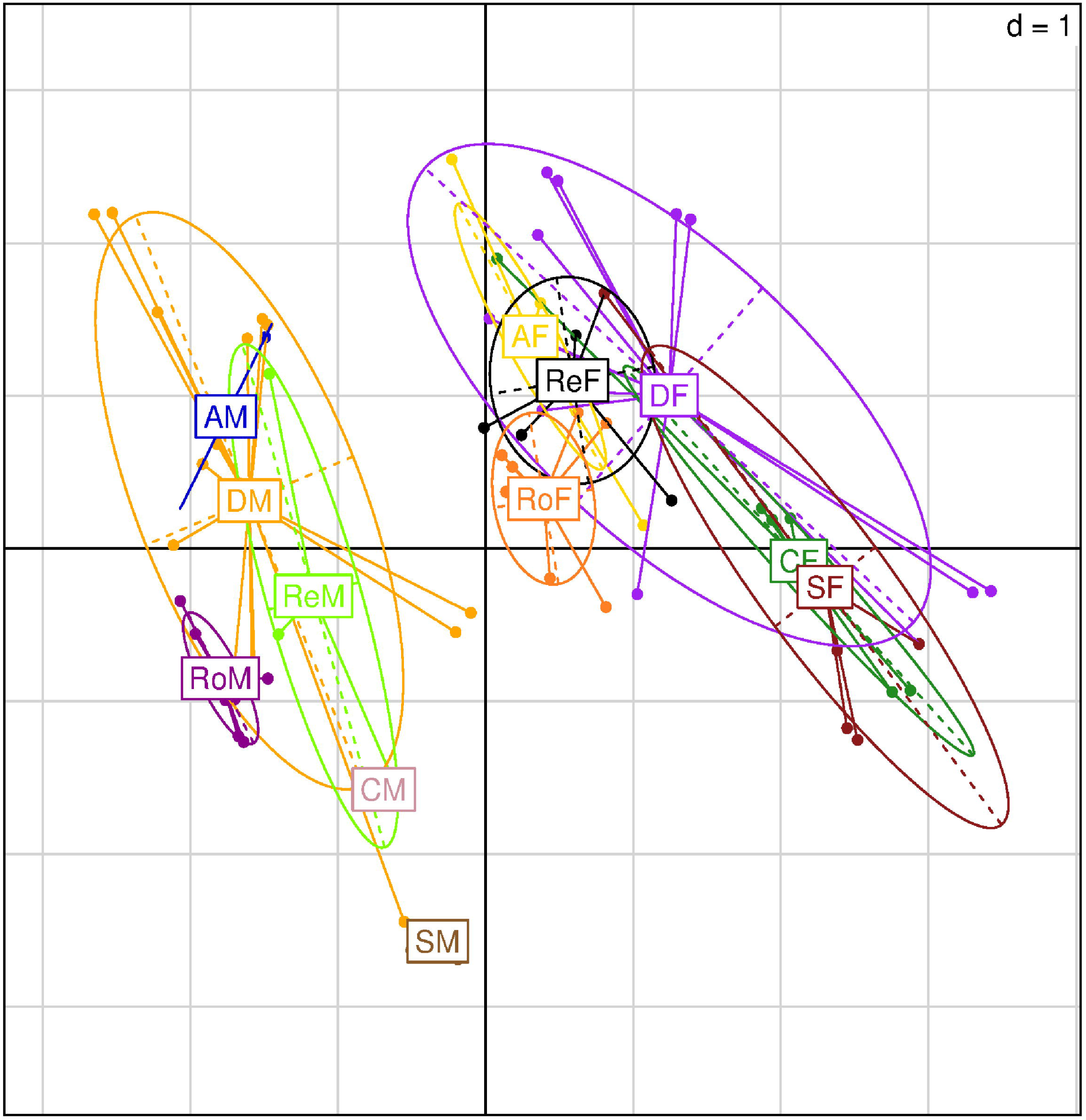
Dispersion of *Phoenix* species in relation to sex on the first biplot of the Principal Component Analysis. **DM:** *P.dactylifera* male; **DF**: *P.dactylifera* female; **CM**: *P.canariensis* male; **CF**: *P.canariensis* female; **SM**: *P.sylvestris* male; **SF**: *P.sylvestris* female; **RoM**: *P.roebelenii* male; **RoF**: *P.roebelenii* female; **AM**: *P.atlantica* male; **Af**: *P.atlantica* female; **ReM**: *P.reclinata* male; **ReF**: *P.reclinata* female.

##### mPdIRDP50 locus

The mPdIRDP50 locus also had Y-linked alleles shared between males of the seven *Phoenix* species (Figure 2). Only *P. roebelenii* had a private allele (mPdIRDP50_177). In Eastern accessions of *P. dactylifera*, we observed Y-linked duplications of the mPdIRDP52 and mPdIRDP50 loci. As in *P. dactylifera, P. sylvestris* and *P. acaulis* displayed duplicated male alleles (Figure 2). The observed identity of mPdIRDP50 allele sizes (Figure 2) seen in eastern accessions of *P. dactylifera, P. sylvestris*, *P. loureroi* and in *P. acaulis* suggests an ancestral origin of these alleles.

##### mPdIRDP80 locus

The mPdIRDP80 locus had a total of seven alleles in the entire sample (Figure 2). The mPdIRDP80_192 and mPdIRDP80_308 alleles were both strictly associated with the male phenotype in the studied species, confirming the linkage to the Y chromosome. The other five alleles (mPdIRDP80_290, mPdIRDP80_292, mPdIRDP80_296, mPdIRDP80_299, mPdIRDP80_312) were shared between males and females and therefore presumed to be X-linked alleles. *P. roebelenii* and *P. acaulis* species had private X alleles, namely the mPdIRDP80_296 allele for *P. roebelenii* and the mPdIRDP80_312 allele for *P. acaulis* (Figure 3).

**Figure 3.**
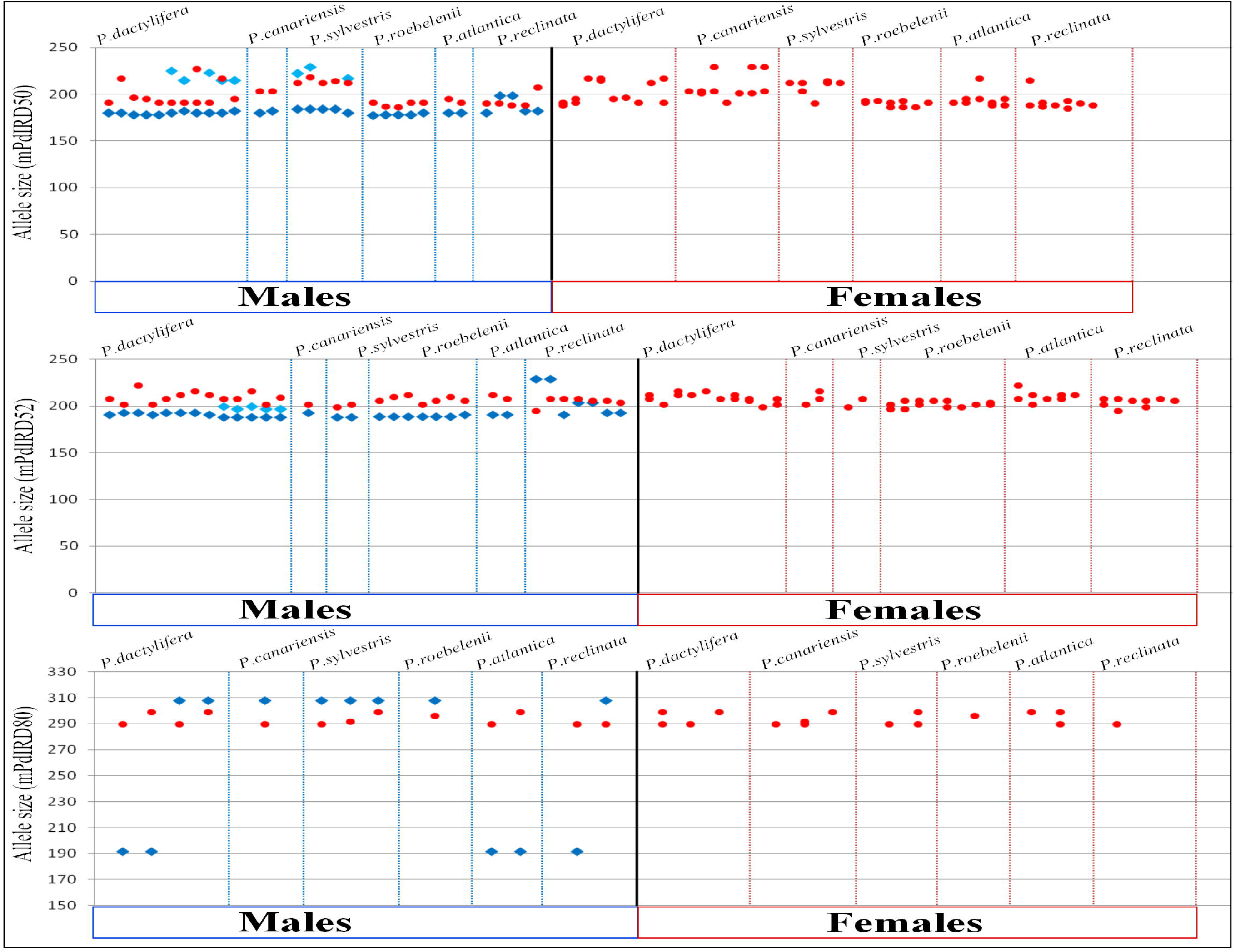
Allelic distribution of mPdIRD50, mPdIRD52 and mPdIRD80 loci within the studied *Phoenix* species. Left side corresponds to male genotypes and right side corresponds to female genotypes. Each dot represents an allele. Alleles shared between male and female individuals (X) are represented in red and male specific alleles (Y) are represented in blue. Duplicated male alleles (Y) are represented in light-blue. Female individuals have only alleles shared between male and female while male individuals have shared alleles and male specific alleles.

The mPdIRDP80 locus showed a high level of conservation among the studied *Phoenix* species, especially for the Y-linked alleles of which only two forms were observed. To further compare X and Y sequences in *Phoenix* spp. and to exclude a possible convergence of SSR allele sizes, we sequenced the mPdIRDP80 locus and performed multiple alignments of X and Y sequences. The alignments of all X sequences showed a shared identity ranging from 95% and 100%. The corresponding western and eastern Y alleles in *P. dactylifera* [20] revealed that the amplified region is identical in the entire male sampling (identity ≥ 99%) (electronic supplementary material, Figure 2, 3 and 4).

Taken together, these results indicate that the sex-linked region bearing the mPdIRDP80, mPdIRDP50 and mPdIRDP52 loci is highly conserved throughout the genus.

Consistent with this conclusion, *R_st_* values [25] (Table 1 and electronic supplementary material, Figure 1), indicate significant genetic differentiation between males and females within each species studied, except for *P. reclinata* (*R_st_* < 0; this value may be biased due to the different numbers of males and females studied, see the electronic supplementary material, Table 4). Moreover, principal component analysis (PCA) yielded separate clusters of males and females (Figure 3). Thus, as for *P*. *dactylifera*, these three loci showed sex linkage in the additional studied species.

#### (ii) Heterozygosity at the sex-linked loci

All males of each *Phoenix* species analysed were heterozygous (Y_allele_/X_allele_) for all three *P. dactylifera* sex-linked microsatellite loci (F*is* ranged from −0.04 to −1, see electronic supplementary material, Table 5), with significant differences from the Hardy–Weinberg genotype frequencies, expected under random mating, F*is* = 0. In contrast, females of each species mostly had significantly positive F*is* values (electronic supplementary material, Table 5), in line with previous reports for X-linked SNPs in date palm females [19].

Overall, these results demonstrate that the XY chromosome system previously observed in *P. dactylifera* is also present in the eight other *Phoenix* species studied here. The findings of this comparative genetic study reveal a common set of orthologous sex-linked markers shared between the additional studied species (*P. canariensis*, *P. atlantica*, *P. reclinata*, *P. sylvestris*, *P. roebelenii*, *P. rupicola*, *P. loureroi* and *P. acaulis*) and *Phoenix dactylifera*.

The Y-linked alleles of the *Phoenix* species studied were assigned to 11 Y haplotypes none of which were found among the 39 X-haplotypes (Figure 4). Therefore, recombination does not occur between the Y and X chromosome regions carrying the mPdIRDP80, mPdIRDP50 and mPdIRDP52 loci in any of the species, strongly suggesting complete sex-linkage of these loci, as in *P*. *dactylifera* [20]. Moreover, the mPdIRDP80 and mPdIRDP50 microsatellite loci are located in the LG12 sex segregating region in the genetic map of date palm corroborating somehow the small size of sex segregating region [21]. Nevertheless, mPdIRDP52 locus was mapped neither in LG12 nor in any other linkage group suggesting that Y region carrying mPdIRDP80, mPdIRDP50 and mPdIRDP52 loci is probably larger than proposed by Mathew *et al*. [21].

**Figure 4.**
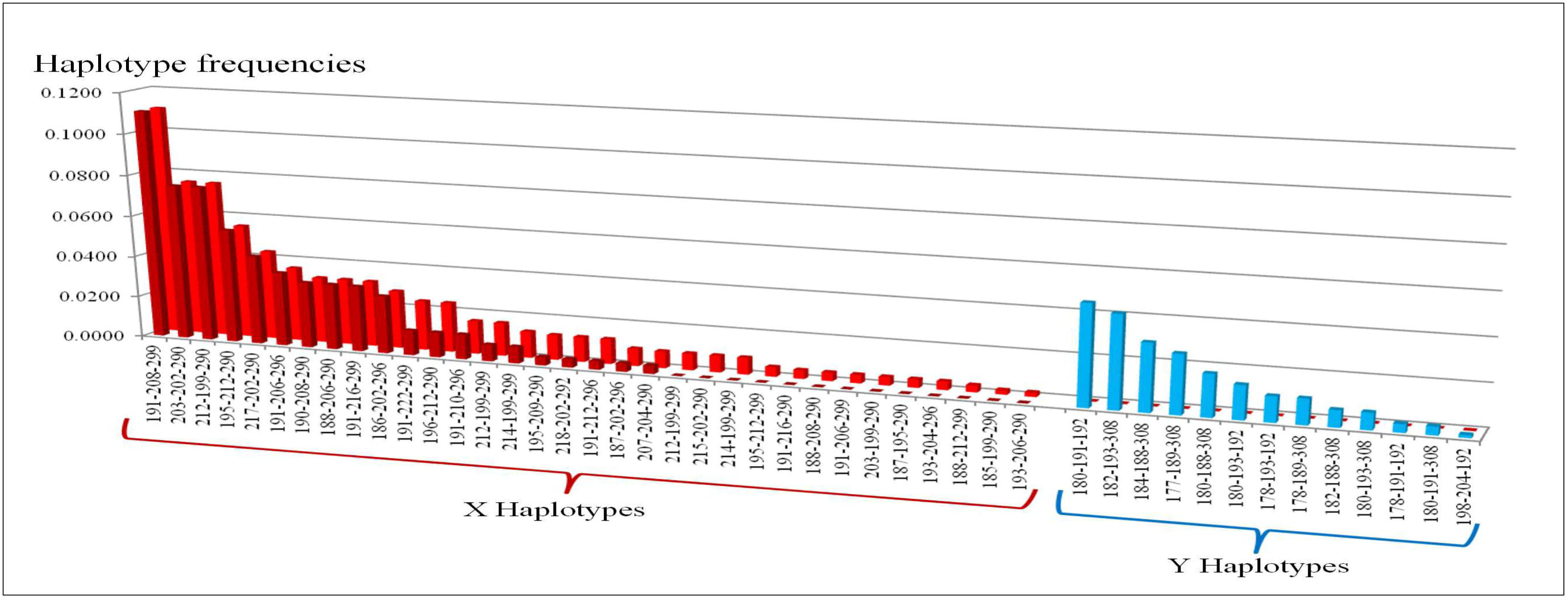
Haplotypes distribution within studied *Phoenix* species analysed as a whole (*P. dactylifera*, *P. reclinata*, *P.sylvestris*, *P. atlantica*, *P. canariensis*, *P. roebelenii* and *P. acaulis*). Light and dark red histograms represent respectively X female and X male haplotypes. Blue histograms correspond to Y haplotypes.

We next investigate whether this sex-linked region has evolved once or several times independently within each species.

### (b) The PdMYB1 gene and the timing of recombination arrest in the *Phoenix* genus

To estimate the time since recombination stopped, estimates of sequence divergence between X and Y alleles are needed (ideally for multiple genes, to test whether different strata exist). The microsatellite loci do not provide such an estimate. One gene is, however, now available. The *PdMYB1* gene (see Methods) is homologous to the predicted *P. dactylifera* transcription factor *MYB86-like* (GenBank accession number XM_008777432.1) and maps in the LG12 (PDK_30s6550963) sex-linked region of the date palm genetic map [21] and the mPdIRDP80 microsatellite locus is localized within its promoter sequence (Figure 5A). We cloned and sequenced the ~ 2 Kb region including the *PdMYB1* gene and mPdIRDP80 (electronic supplementary material, Figure 5). The *PdMYB1* has three exons of 135, 128 and 963 pb, and two introns of 191 and 102 pb, and the mPdIRDP80 sequence is 176 pb upstream of the first exon (Figure 5 A). Sequencing of the mPdIRDP80 locus confirmed the observed sizes of X and Y alleles from capillary gel genotyping (Figure 5 A). We cloned the *PdMYB1* gene in the seven related *Phoenix*, and identified their X/Y pairs based on the sizes of the mPdIRDP80 X and Y alleles. All males had different X and Y sequences while all females had only the X sequences.

**Figure 5.**
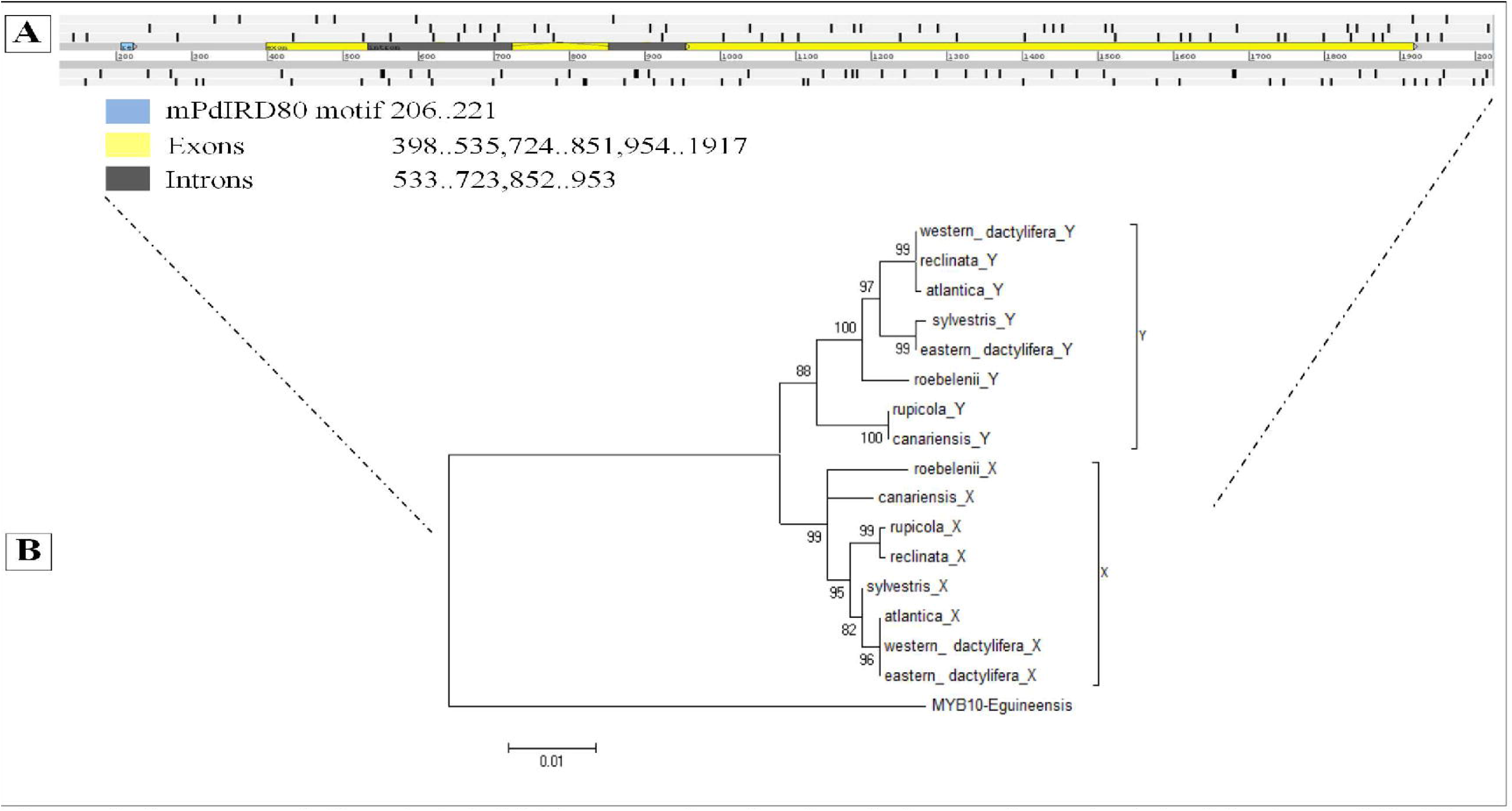
Structure of *Phoenix PdMYB1* gene and molecular phylogenetic analysis by Maximum Likelihood method. **A.** The structure of the *PdMYB1* gene was predicted based on multiple alignments including the *P.dactylifera* transcription factor *MYB*86-like and the orthologous sequences of other species. Blue, yellow and grey rectangles indicate respectively the position of mPdIRDP80 motif, exons and introns of the *PdMYB1* gene. **B.** The tree with the highest log likelihood (-3368.6438) is shown. The percentage of trees in which the associated taxa clustered together is shown next to the branches (1000 replications). Initial tree (s) for the heuristic search were obtained automatically by applying Neighbor-Join and BioNJ algorithms to a matrix of pairwise distances estimated using the Maximum Composite Likelihood (MCL) approach, and then selecting the topology with superior log likelihood value. The tree is drawn to scale, with branch lengths measured in the number of substitutions per site. The analysis involved 19 nucleotide sequences. All positions with less than 95% site coverage were eliminated. That is, fewer than 5% alignment gaps, missing data, and ambiguous bases were allowed at any position. There were a total of 1494 positions in the final dataset. Corresponding orthologous of the *Elaeis guineensis PdMYB1* gene was used as the outgroup. X and Y represent the X- and Y-specific alleles, respectively.

We reconstructed the phylogeny of the X and Y copies of the *PdMYB1*gene from the newly studied *Phoenix* species, and found distinct Y and X alleles clusters, rather than a clustering by species of origin (Figure 5 B). The same result was also obtained with the distribution of X and Y mPdIRDP80, mPdIRDP50 and mPdIRDP52 SSRs alleles among the *Phoenix* species by PCA (see the electronic supplementary material, Figure 6). These results provide evidence that recombination between X and Y alleles of the *PdMYB1*gene and the sex-linked loci stopped before the *Phoenix* spp. split. Similarly, phylogenetic analysis suggested that recombination stopped before the speciation of the dioecious species *S. latifolia*, *S. dioica*, and *S. diclinis*, whereas sequences of the SIX1/SIY1 gene may have undergone suppressed recombination independently in these species [3].

**Figure 6.**
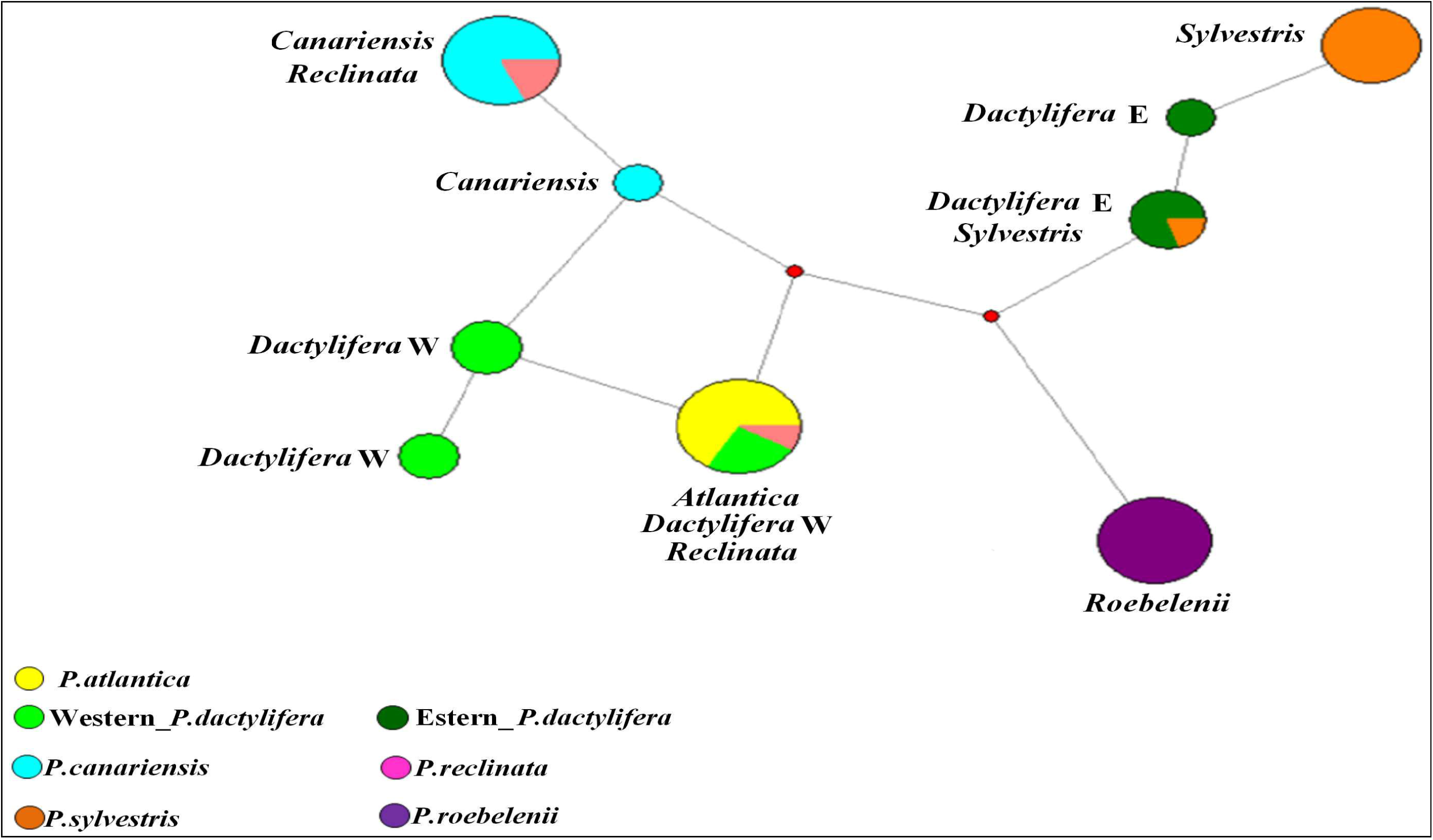
Date palm Y-SSR haplotype Network. Red dots indicate the median vectors corresponding to the hypothesised haplotypes required to connect the existing ones.

### (c) Y chromosome evolution in the genus *Phoenix*

The network based on the Y haplotypes revealed at least two major Y chromosome lineages (Figure 6). The first includes four species, two from the Atlantic islands (*P. canariensis* and *P. atlantica*), one from sub-Saharan Africa (*P. reclinata*) plus the western *P*. *dactylifera* lineage (North Africa). The second lineage includes three species, the eastern *P. dactylifera* lineage, from Pakistan and the Middle East, and two species from Asia (*P. sylvestris and P. roebelenii*) (Figure 6). In the eastern lineage, the *P. roebelenii* group appears divergent from the *P. sylvestris* and eastern *P. dactylifera* groups and has private Y haplotypes. The two male lineages within *P. dactylifera* were previously identified [20]. The observation that these haplotypes are also found in other *Phoenix* species suggests a split into two male paternal lineages from an ancestral Y population before speciation, with the two Y-haplotypes transmitted to the evolving species and becoming fixed in different species. However, the presence of haplotypes within *P. dactylifera* is surprising. It could be due to ancestral polymorphism maintained on the Y chromosome or to more recent introgressions but this still need to be investigated.

## 4. Conclusion

According to fossils of *Phoenix* male flowers, dioecy could have emerged in the genus during the Eocene (~ 49.5 mya) [10, 38].

Our data bring evidence that recombination arrest evolved prior to speciation in the genus *Phoenix*. Furthermore, the entire extant *Phoenix* species are dioecious which allows to conclude that sex chromosomes and dioecy evolved only once in this lineage. The time at which recombination stopped in the *Phoeniceae* can be estimated by analyzing X/Y divergence of other sex-linked genes especially from the non-recombining regions of the date palm sex-linkage group [21].

## Data accessibility

*The datasets supporting this article have been uploaded as part of the Supplementary Material.*

## Competing interests

*‘We have no competing interests.’*

## Author’s contributions

FAB conceived and planned the project, EC and KC, performed the in silico research of SSR, EC, NC, JCP performed the DNA preparations, EC performed the genotyping, AC performed the cloning and sequencing of the MYB gene, EC performed the genetic and statistical analyses, EC, SZ, AC, JCP, SS, SG, ASH and FAB, discussed the results, EC and FAB wrote the paper. All authors gave final approval for publication.

## Acknowledgements

We thank Ali Zouba, Adbelmadjid Rhouma, Karim Kadri, Ahmed Othmani (CRRAO, Tunisia), Abdourahman Daher, Sabira Abdoulkader (CERD, Djibouti),, Claudio Littardi (CRSP, Italy), Jose Plumed (Botanical garden of Valencia, Spain), Marco Ballardini (CRAFSO, Italy), Muriel Gros-Balthazar (IRD, France), Robert Krueger (USDA, USA), Emmanuel Spick (Jardin des Plantes de Montpellier, France), Bill Baker (Kew Gardens, UK) and Alain Rival (Cirad, France) for providing plant material; Sylvain Santoni and Muriel Latreille (INRA, France) for technical assistance; and James Tregear for corrections to the manuscript.

## Funding

This work was supported by the AUF-Mersi Project, the ‘Ministère de l’Enseignement Supérieur et de la Recherche’ of Tunisia and the Qatar National Research Fund (NPRP-EP X-014–4–001).

